# Rad51 catalytic mutants differentially affect the Rad51 nucleoprotein filament *in vivo*

**DOI:** 10.1101/070276

**Authors:** Maureen M. Mundia, Alissa C. Magwood, Mark D. Baker

**Author notes:** **Correspondence:** Dr. Mark D. Baker, Dept of Molecular and Cellular Biology, College of Biological Sciences, University of Guelph 50 Stone Road East, Guelph, Ontario N1G 2W1 Canada.

## Abstract

In this study, we utilized mouse hybridoma cell lines stably expressing ectopic wild-type Rad51, or the Rad51-K133A and Rad51-K133R catalytic mutants deficient in ATP binding and ATP hydrolysis, respectively, to investigate effects on the Rad51 nucleoprotein filament *in vivo*. Immunoprecipitation studies reveal interactions between ectopic wild-type Rad51, Rad51-K133A and Rad51-K133R and endogenous Rad51, Brca2 and p53 proteins. Importantly, the expression of Rad51-K133A and Rad51-K133R catalytic mutants (but not wild-type Rad51) targets endogenous Rad51, Brca2 and p53 proteins for proteasome-mediated degradation. Expression of Rad51-K133R significantly reduces nascent DNA synthesis (3’ polymerization) during homologous recombination (HR), but the effects of Rad51-K133A on 3’ polymerization are considerably more severe. Provision of additional wild-type Rad51 in cell lines expressing Rad51-K133A or Rad51-K133R does not restore diminished levels of endogenous Brca2, Rad51 or p53, nor restore the deficiency in 3’ polymerization. Cells expressing Rad51-K133A are also significantly reduced in their capacity to drive strand exchange through regions of heterology. Our results reveal an interesting mechanistic dichotomy in the way mutant Rad51-K133A and Rad51-K133R proteins influence 3’ polymerization and provide novel insight into the mechanism of their dominant-negative phenotypes.

## INTRODUCTION

Homologous recombination (HR) is a complex multistep process for high-fidelity repair of DNA double-strand breaks and the recovery of stalled or collapsed DNA replication forks. Consequently, HR plays a pivotal role in the control of genome stability and tumor suppression. At the heart of the HR process is the pre-synaptic filament, which is formed when single-stranded DNA (ssDNA) generated by nucleolytic DNA end-resection is coated with DNA recombinase. DNA recombinases comprise an evolutionarily-conserved family typified by the *Escherichia coli* RecA protein and include additional members such as bacteriophage T4 UvsX, *Saccharomyces cerevisiae* Rad51, and mammalian Rad51 proteins.^1^ The DNA recombinase-coated nucleoprotein filament is essential for the early homology search and strand invasion steps of HR. Strand invasion pairs the broken strand of DNA with its unbroken complementary partner, generating a synaptic intermediate that primes new DNA synthesis (3’ polymerization) thereby replacing nucleotides originally lost at the DNA break site. Subsequent steps depend on the particular HR pathway and may include dissolution of the strand containing the newly-synthesized DNA, generating a non-crossover product, or maturation to a double-Holliday junction intermediate that can be cleaved to generate either crossover or non-crossover products.^2^

The assembly of recombinase enzymes onto ssDNA requires recombination mediator proteins to remove firmly-bound ssDNA binding proteins, which pose a barrier to pre-synaptic filament formation.^3^ In mammalian cells, the breast cancer, type 2 susceptibility (BRCA2) protein fulfills a recombination mediator function specifically directing Rad51 filament assembly on replication protein A (RPA)-coated ssDNA.^4–6^ BRCA2 interacts with RAD51 via a series of eight BRC repeats and a C-terminal binding site. BRCA2 directs loading of the ATP-bound form of RAD51 onto ssDNA and by decreasing the rate of ATP hydrolysis, promotes assembly of the active ATP-bound form of RAD51 in the nucleoprotein filament. BRCA2 also stabilizes the ADP-bound form of RAD51 permitting exchange of ADP for ATP, thus further increasing the content of ATP-bound RAD51 in the nucleoprotein filament that is active in HR. Therefore, the ATP-bound form of RAD51 is required for effective RAD51 binding to ssDNA, while ATP hydrolysis is considered important for RAD51 turnover. The dynamic state of the Rad51 nucleoprotein filament ensures RAD51 nucleoprotein filament growth, and the subsequent homology search and strand invasion steps of HR.^7^

Hallmarks of recombinase enzymes are conserved Walker A and B motifs that function in ATP binding. Substitution of a conserved lysine in the Walker A motif (K133 in mammalian RAD51) severely affects ATP binding/hydrolysis: mammalian RAD51-K133A and RAD51-K133R are defective in ATP binding and hydrolysis, respectively ^8–10^, and are similar to the ATP-hydrolysis defective *Saccharomyces cerevisiae* dominant-negative mutants, Rad51-K191A and Rad51-K191R mutants, respectively.^11–13^ We have shown previously^14^ that mouse hybridoma cell lines stably expressing mouse Rad51-K133R or Rad51-K133A display sensitivity to DNA damaging agents, defects in gene targeting, and abnormal intrachromosomal homologous recombination. Expression of human RAD51-K133R in mouse ES cells causes hypersensitivity to DNA damaging agents, decreased rates of spontaneous sister chromatid exchange, and a reduction in DSB-induced recombination between homologous repeats.^15–16^ In addition, expression of the Rad51 catalytic mutants impedes anaphase chromosome segregation^17^, DNA replication re-start, and the maintenance of chromosomal stability^18^. In all cases, the attained expression levels of the mutant Rad51 alleles, especially K133A, are lower than the corresponding wild-type protein^14,15,18^, suggesting that even low levels of catalytic Rad51 mutants are toxic.

The mechanism whereby the Rad51-K133A and Rad51-K133R mutants affect HR is not known although the possibility that homotypic interactions with endogenous wild-type Rad51 and formation of a mixed Rad51 nucleoprotein filament with aberrant function has been suggested.^15,18^ The effect of catalytic Rad51 mutant alleles on Rad51 nucleoprotein filament formation and function *in vivo* has not been validated and is the subject of the following investigation.

## MATERIALS AND METHODS

### Cell lines and culture conditions

The origin of the mouse hybridoma cell line igm482 and derivatives expressing FLAG-tagged wild-type Rad51, or Rad51-K133A and Rad51-K133R catalytic mutants, and the conditions used for hybridoma cell culture in Dulbecco’s modified Eagle’s medium (DMEM) have been described elsewhere.^14,19,20^ When required for transformant culture, DMEM was supplemented with the appropriate selectable agents, G418 (600 μg/mL), puromycin (7.5 μg/mL) or hygromycin (700 μg/mL). When required, hybridoma cells (in DMEM) were exposed to a 10 μM solution of the proteasome inhibitor MG132 dissolved in dimethyl sulfoxide (DMSO)^21^ (CalBiochem). As a control for effects of the solvent on protein expression, cells were exposed to 0.1% DMSO (in DMEM) for 60 min. When required, cells were treated with 60 μg/mL cycloheximide (Sigma-Aldrich). Cell viability and density were determined by Trypan Blue stain exclusion (HyClone) and a hemocytometer (Bright-Line).

### Plasmids, DNA transfer and transformant recovery

Plasmid DNA was propagated in *Escherichia coli* (DH5α) and extracted using the PureLink™ HiPure plasmid maxiprep kit (Life Technologies). Restriction enzymes were purchased from New England BioLabs (Mississauga, Ontario) and used in accordance with the manufacturer’s instructions. Derivatives of the p3X-FLAG-CMV^™^-10 (Sigma) expression vector bearing 3X N-terminal FLAG-tagged wild-type mouse Rad51 were constructed and verified as reported previously^14^ and used for expression of wild-type Rad51 in transformants expressing Rad51-K133A or Rad51-K133R. The plasmid pTΔCμ_858/2290_ used in the detection of nascent DNA synthesis during the early steps of mammalian homologous recombination was described previously^22,23^ and contains 858 bp and 2290 bp left and right arms of homology, respectively, to the haploid hybridoma chromosomal immunoglobulin μ locus. The plasmid, pTΔCμ_858*/2290_ is a pTΔCμ_858/2290_ derivative in which otherwise continuous homology in the left vector arm is interrupted by a 4-bp insertion positioned 211 bp from the vector 3’ end. The 4-bp insertion converts a genomic *SacI* site to a vector-borne *EcoRV* site.^24^ Transfer of plasmid DNA to hybridomas was performed by electroporation^22^ and individual transformants were recovered by limited dilution cloning.^14^

### RT-PCR

Total RNA was prepared by TRIzol extraction (Life Technologies). The RNA was then treated with DNaseI (Roche) and reverse transcribed using Superscript II (Life Technologies). For RT-PCR analysis, the PCR reactions were first optimized to determine the optimal cycle number in which amplification and detection of the transcripts was within linear range. A total of 600 ng of cDNA was used for each RT-PCR reaction. Primers specific for RT-PCR analysis of *Brca2, p53,* and the control single-copy chromosomal immunoglobulin *μ* gene have been described previously.^25,26^ For analysis of total Rad51 transcript levels, we utilized forward (5’-GCTTGAAGCAAGTGCAGA-3’) and reverse (5’-CATAGCTTCAGCTTCAGG-3’) primers, which amplify a 960 bp fragment in both the FLAG-tagged and endogenous Rad51 cDNA. The forward primer 5’-CAAGGATGACGATGACAAGC-3’ and the aforementioned reverse primer were used to specifically amplify a 1079 bp fragment from the FLAG-tagged Rad51 cDNA.

### Protein analysis

Western blot and immunoprecipitation analyses were performed according to Magwood *et al*.^25^ with the exception that rabbit IgG serum (#31235, ThermoScientific) was used as a non-specific antisera in immunoprecipitation. The following primary antibodies were used for Western blot and/or immunoprecipitation analyses; anti-human Rad51 (14B4, Abcam), antihuman Rad51 (H92, Santa Cruz Biotechnology), anti-FLAG (M2, Sigma-Aldrich), anti-β-actin (AC-15, Sigma-Aldrich), anti-human histone H1 (A-E4, Santa Cruz Biotechnology), anti-human p53 (FL-393, Santa Cruz Biotechnology), anti-human BRCA2 (Ab27976, Abcam), and anti-ubiquitin (07-2130, EMD Millipore). Immunoblot signals were detected with ECL-Prime reagent (GE Healthcare) using the appropriate HRP-coupled goat anti-mouse IgG (1030-05, Southern Biotech), goat anti-rabbit IgG (111-036-047, Jackson Immunoresearch) or Veriblot for IP (Ab131366, Abcam) secondary antibodies.

### Measurement of 3’ polymerization during homologous recombination

The formation of nascent DNA that follows the strand invasion event of homologous recombination (3’ polymerization) was measured by a sensitive PCR assay as described previously.^22^

### Data analysis

The Gel Doc system (BioRad) and Quantity One imaging software version 4.4.6 (BioRad) were utilized for densitometric analysis of band intensity when determining the intensity of specific PCR bands in the 3’ extension assay as well as for Western blot and RT-PCR analyses. Statistical analysis was performed by one-way analysis of variance (ANOVA) and Tukey’s-HSD (Honestly Significant Difference) test, or *t*-test using VassarStats statistics software available online (http://vassarstats.net/). Significance was assessed at *p* ≤ 0.05. Error bars represent the standard error of the mean.

## RESULTS

### Rad51 catalytic mutants feature reduced levels of endogenous Rad51, Brca2 and p53 proteins

Derivatives of the hybridoma cell line, igm482^19,20^ stably expressing N-terminal 3X-FLAG-tagged wild-type Rad51 or the FLAG-tagged ATP-catalytic site mutants, Rad51-K133A and Rad51-K133R, that are deficient in ATP binding and ATP hydrolysis, respectively,^8–10, 12^ have been described previously.^14^ The ~42 kDa FLAG-tagged Rad51 and 37 kDa endogenous Rad51 proteins were identified by Western blot analysis of whole cell extracts (Fig. 1a). Interestingly, the ~42 kDa Rad51-K133R (lanes 3 and 4) and Rad51-K133A (lanes 7 and 8) mutant proteins are expressed at considerably lower levels than wild-type FLAG-Rad51 in the isogenic WT5 cell line (lane 1) or the wild-type FLAG-Rad51 that is co-expressed in derivatives of K133R-11 (lane 5) and K133A-6 (lane 9). The Rad51-K133A expression, in particular, is at such low levels that the FLAG-tagged mutant protein is not visible in Western blot analysis using the anti-Rad51 antibody (14B4). We can, however, detect the protein variant using the anti-FLAG antibody (M2) which recognizes the three N-terminal FLAG epitopes in the FLAG-tagged Rad51 protein, suggesting that the three epitopes provide the opportunity for an amplified signal that allows detection of the protein even at very low quantities. Immunoblot analysis using the anti-FLAG antibody followed by densitometric analysis determined that the transgene/β-actin ratios in cell lines expressing wild-type FLAG-Rad51 were ~1.6 and higher, while transgene/β-actin ratios ranged between 0.7-1.0 in cell lines expressing Rad51-K133R, and ~0.3 and lower in cell lines expressing Rad51-K133A. The low expression attained suggests toxicity associated with recovering transformants expressing the Rad51 catalytic mutants. Indeed, we previously showed that even low expression of the Rad51-K133A allele reduced the doubling time of hybridoma cells^14^, a result consistent with other studies suggesting toxic effects arising from over-expression of Rad51 catalytic mutants.^15–18^

**Fig. 1.**
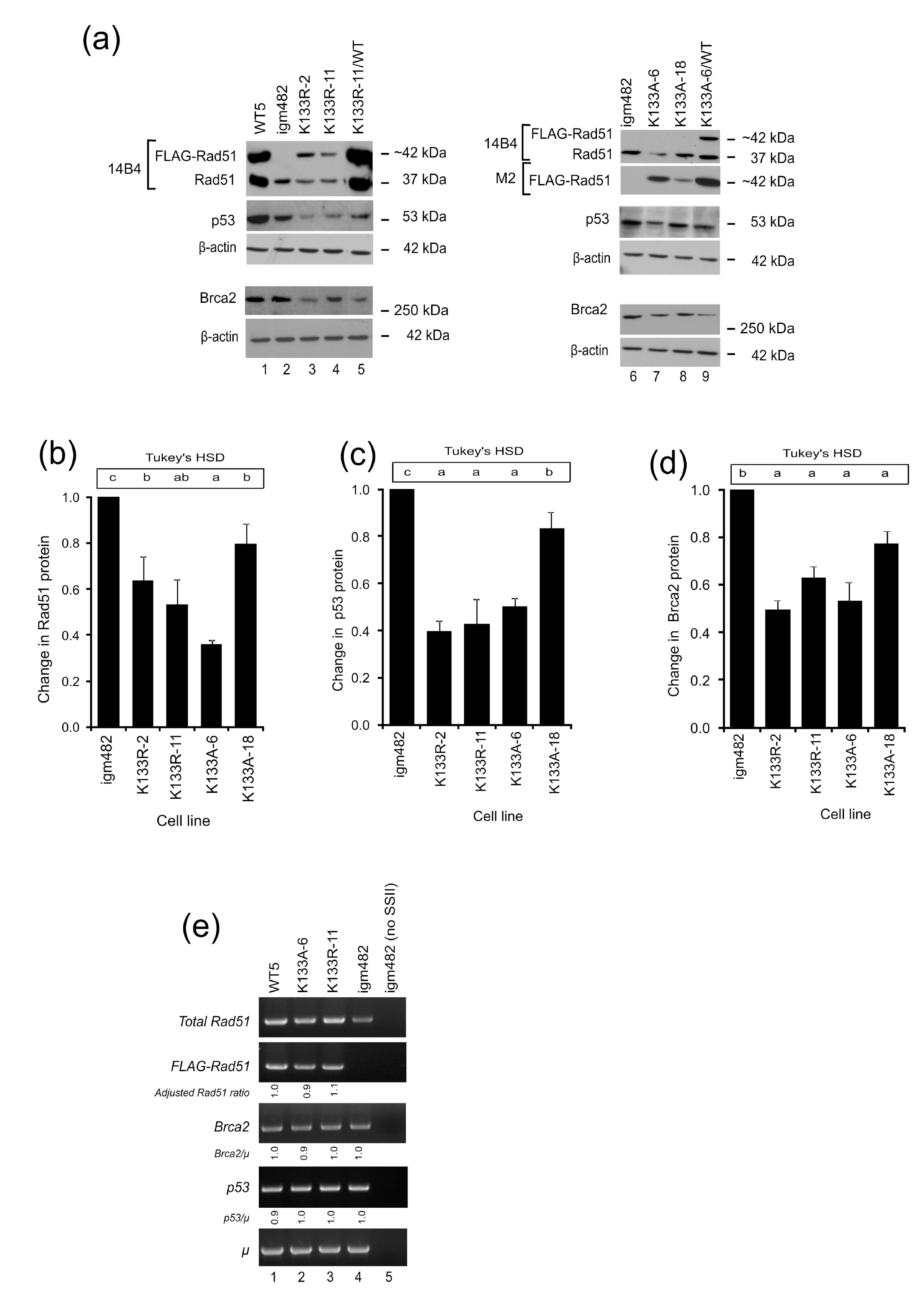
Rad51, p53 and Brca2 levels. (a) Representative Western blots of Rad51 (14B4), FLAG-Rad51 (M2), p53 (FL-393), and Brca2 (Ab27976) proteins. β-actin (AC-15) is presented as a control for protein loading. (b-d) Fold change in endogenous proteins (Rad51, p53, and Brca2) compared to control igm482 cells ± standard error of the mean as determined by densitometric analysis of Western immunoblots from at least 3 independent experiments. One-way ANOVA and Tukey’s-HSD analysis were used to assess statistically significant differences between cell lines. Means indicated by the same lower-case letter are not statistically different (p≤0.05). (e) RT-PCR analysis of Rad51, Brca2 and p53 transcript levels. Densitometric analysis of band intensity was used to determine the various transcript levels relative to that of the chromosomal immunoglobulin *μ* gene. The Brca2/μ and p53/μ ratios were determined relative to control igm482. The calculation for the Adjusted Rad51 ratio is explained in the text (see Results section). As a control for cDNA synthesis, SuperScript II reverse transcriptase (SSII) was omitted from the sample in lane 5.

Magwood *et al.*^26^ reported that endogenous levels of Rad51 and Brca2 depend on each other and that p53 levels adjust accordingly to accommodate the status of the other two proteins. Subsequently, we investigated the status of these proteins in cell lines stably expressing wild-type FLAG-Rad51, and the Rad51-K133A and Rad51-K133R catalytic mutants. Quite unexpectedly, we observed that the level of endogenous Rad51 protein is reduced in cell lines expressing moderate levels of Rad51-K133R (lanes 3 and 4) and Rad51-K133A (lane 7) compared to that in control igm482 cells (lanes 2 and 6). The reduction in endogenous Rad51 is less dramatic in the cell line expressing very low amounts of the transgene (K133A-18; lane 8). Densitometric analysis of Western blots from three independent experiments revealed that the Rad51 catalytic mutants have significantly reduced endogenous Rad51 levels; K133R-2 and K133R-11 are ~40% reduced, while K133A-6 and K133A-18 are ~60% and ~20% reduced, respectively (Fig. 1b).

Our igm482 hybridoma expresses a wild-type *p53* gene.^26^ Therefore, whole cell extracts were also analyzed for total cellular wild-type p53 protein. As shown in Fig. 1a, in contrast to the control igm482 hybridoma (lanes 2 and 6), hybridoma cell lines expressing moderate levels of Rad51-K133R (lanes 3 and 4) and Rad51-K133A (lane 7) feature lower levels of p53 protein. Cell line #18 expressing a lower level of the Rad51-K133A catalytic mutant shows a slight reduction in endogenous p53 protein (lane 8). Endogenous p53 protein is elevated by wild-type FLAG-Rad51 expression perhaps as a result of protection through self-interaction^27^ (cell line WT5; lane 1) and, notably, expression of wild-type Rad51 in cell lines K133R-11/WT (lane 5) and K133A-6/WT (lane 9) cannot restore normal levels of p53. Replicate experiments revealed that the three cell lines K133R-2, K133R-11, K133A-6 show ~50-60% reduction in endogenous p53 and K133A-18 is ~15% reduced (Fig. 1c).

Western blot analysis of whole cell extracts using anti-BRCA2 antibody (Ab27976) (Fig. 1a) revealed that in comparison to control igm482 (lanes 2 and 6) and WT5 (lane 1) hybridoma cells, the levels of endogenous Brca2 protein are reduced in cell lines expressing Rad51-K133R (lanes 3 and 4) and Rad51-K133A (lanes 7 and 8). Notably, expression of wild-type FLAG-Rad51 cannot restore endogenous Brca2 levels in cell lines co-expressing the Rad51 catalytic mutants, K133R-11/WT (lane 5) and cell line K133A-6/WT (lane 9). Replicate experiments revealed that the level of Brca2 is reduced in cell line K133A-18 by ~10-20%, while the three moderately-expressing cell lines K133R-2, K133R-11, K133A-6 show ~40-50% reduction (Fig 1d). Thus, we suggest that expression of the Rad51 catalytic mutant triggers depletion of endogenous Rad51/Brca2, which in turn leads to a decrease in p53.

As shown in Fig. 1a, a higher level of “endogenous Rad51” protein is noticeable in cell lines WT5, as well as K133R-11/WT and K133A-6/WT co-expressing wild-type FLAG-Rad51, compared to control igm482 cells. The augmented levels of “endogenous” Rad51 protein in cell lines over-expressing Rad51 proteins tagged with N-terminal 3X-FLAG may stem from three inframe ATG (AUG) start codons in the mouse *Rad51* cDNA and the subsequent opportunities for multiple internal translation initiations^26^. To test this, we generated cell lines expressing N-terminally FLAG-tagged Rad51 in which the three in-frame start codons were changed to AGG (arginine) such that the expression of the cDNA would only be from the FLAG start codon (Fig. S1a). In mammalian cells, ribosomes can initiate translation at several non-AUG codons with varying efficiency, with the exception of AGG and AAG.^28^ We observed that the cell lines expressing the N-terminal Rad51 variants feature endogenous Rad51 levels that are similar to control igm482 (Fig. S1b top panel, compare lane 2 with lanes 3-5). Thus, we believe that the augmented level of “endogenous” Rad51 that we observe in our FLAG-tagged Rad51 expressors is indeed a result of the three ATG codons in wild-type mouse Rad51 cDNA. Evidently, caution needs to be exercised when interpreting levels of “endogenous” Rad51. However, because FLAG-Rad51 expression levels are quite low in the catalytic mutants, we imagine that the contribution to the “endogenous” Rad51 in these cell lines would be minimal.

RT-PCR was used to determine whether the reduction in endogenous Rad51, Brca2, and p53 proteins resulted from lowered mRNA transcripts as referenced against the single-copy chromosomal immunoglobulin *μ* gene in the hybridoma cells (Fig. 1e). Depleted levels of endogenous Rad51 protein in Rad51-K133A and Rad51-K133R were examined using internal Rad51 primers (that detect all Rad51 transcripts, both FLAG-tagged and “endogenous”) and a pair of primers specific for FLAG-Rad51. To examine any differences in endogenous Rad51 transcript levels, we subtracted the contribution of a normal level of endogenous Rad51 (in igm482) (panel 1, lane 4) from the expression of WT5 (panel 1, lane 1), K133A-6 (panel 1, lane 2) and K133R-11 (panel 1, lane 3), and then divided by the expression of FLAG-Rad51 in these cell lines (panel 2, lanes 1-3). This calculation is expressed as: [(total Rad51/μ) – (igm482 Rad51/μ) / (FLAG/μ)] and represented any change in endogenous expression as caused by overexpression of the respective transgenes. When standardized to the expression of the wild-type transgene in WT5 (denoted as Adjusted Rad51 ratio), it was evident that expression of Rad51-K133A and Rad51-K133R does not affect the levels of endogenous Rad51 transcripts. The similar *Brca2/μ* (panel 3) and *p53/μ* (panel 4) ratios also suggest that changes in mRNA do not account for the reduction in Brca2 and p53 proteins in cell lines expressing Rad51-K133A and Rad51-K133R, or the increase in Brca2 and p53 protein in cell line WT5 expressing wild-type Rad51 (Fig. 1a).

### Depletion of endogenous Rad51, Brca2 and p53 in Rad51 catalytic mutants is due to proteasome-mediated degradation

Previously, we showed that proteasome-mediated degradation can maintain a balanced equilibrium in the cellular levels of endogenous wild-type Rad51, Brca2 and p53 proteins.^26^ In this study, we were interested in determining whether the decreased levels of endogenous Rad51, Brca2 and p53 proteins resulting from expression of the Rad51 catalytic mutants, K133A and K133R, were due to proteasome-mediated degradation.

First, the intrinsic stability of the proteins was examined in half-life studies using cycloheximide (CHX), a well-known inhibitor of protein biosynthesis in eukaryotic cells.^29^ The term “half-life” is used here to refer to the time point after treatment with CHX, at which the amount of protein is half that present in cells that have not been treated (0 h). A time course treatment of igm482 cells (Fig. 2a, panel 1) with 60 μg/ml CHX revealed a gradual decrease in the endogenous levels of Rad51 (lanes 1-5), Brca2 (lanes 6-10) and p53 ( lanes 11-15). The levels of β-actin remain unaffected: the half-life of β-actin is more than 48 h.^30,31^’ In contrast, in the K133R-11 (panel 2, lanes 1-15) and K133A-6 (panel 3, lanes 1-15) cell lines, the corresponding protein levels drastically decline between the 2 h and 4 h treatments and little to no protein was observed after 4 h. We then performed densitometric analyses of Western blots from three independent experiments and calculated the average fold reduction of Rad51, Brca2, and p53 at different time points relative to the amount of protein in the untreated cells to obtain the kinetic analyses shown in Fig. 2b-d. As summarized in (Table 1, the results show that expression of the catalytic mutants significantly increases the susceptibility of the endogenous Rad51, Brca2 and p53 proteins to degradation.

**Fig. 2.**
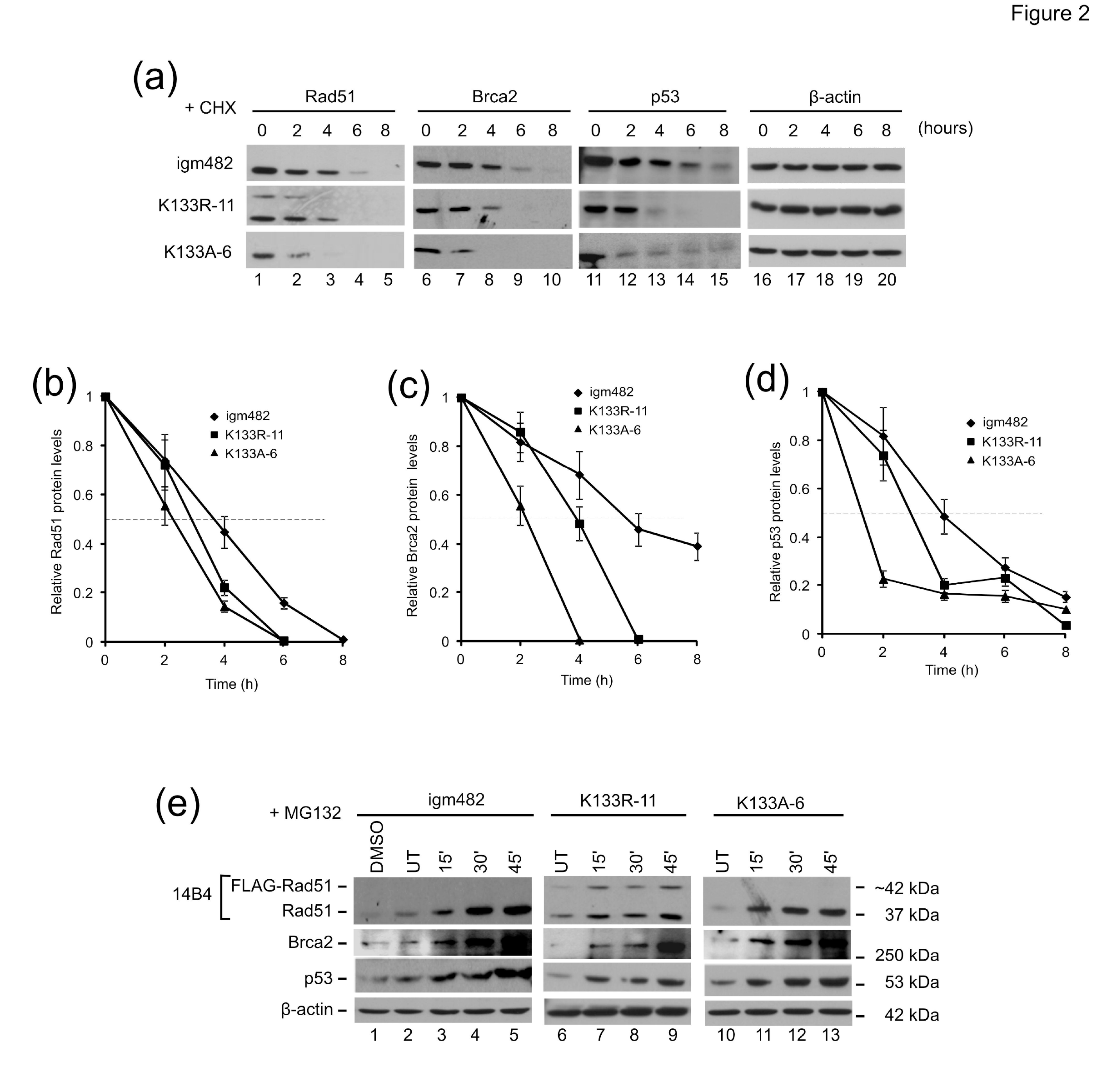
Rad51, p53 and Brca2 protein stability. (a) Analysis of the effects of the protein synthesis inhibitor cycloheximide (CHX) on endogenous Rad51, Brca2, and p53. Time of exposure (hours) to 60μg/ml CHX is indicated. (b-d) Analysis of protein half-life for Rad51, Brca2 and p53. Protein levels were determined by densitometric analysis and the level of protein at each time point relative to untreated (UT) ± standard error of the mean is shown. (e) Analysis of the effects of the proteasome inhibitor MG132 on endogenous Rad51, Brca2, and p53 protein levels. Time of exposure (minutes) to 10 μM MG132 is indicated. Cells were also exposed to 0.1% dimethyl sulfoxide (DMSO) (the solvent for MG132) for 60 min as a control (lane 1). β-actin is presented as a control for protein loading.

**Table 1.**
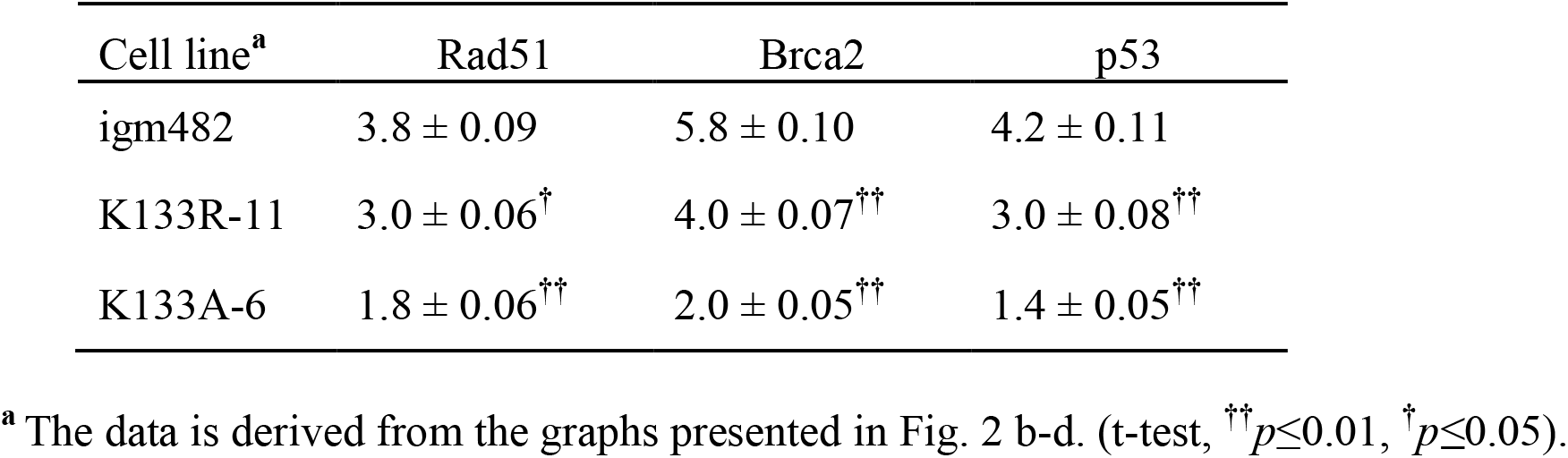
Estimated protein half-lives from cycloheximide treatment experiments

Next, time-course treatment with 10 μM MG132, a proteasome inhibitor^21^, revealed a consistent increase in the endogenous levels of Rad51, Brca2 and p53 proteins in igm482 cells over the 45 min treatment highlighting normal proteasome turnover (Fig. 2e, panels 1-3: lanes 2-5). In cell lines expressing the FLAG-tagged Rad51-K133R and Rad51-K133A catalytic mutants, the recovery in the levels of these proteins appeared more rapid, often as early as 15 min following MG132 exposure (panels 1-3: compare lane 6 with 7, and lane 10 with 11). The levels of β-actin are unaffected by 10 μM MG132 treatment consistent with previous studies^32,33^. The fold increase of endogenous Rad51, p53 and Brca2 proteins following 15 min MG132 exposure compared to the untreated samples was determined by densitometric analysis of three independent experiments and the fold changes are presented in (Table 2. Indeed, an enhanced recovery of the Rad51 and p53 proteins was observed in cell lines expressing the Rad51 catalytic mutants. However, the increase in Brca2 was not as dramatic, perhaps owing to the large size of this protein. Overall, the above results suggest that Rad51 catalytic mutants elevate the normal rate of proteasome-mediated turnover of endogenous Rad51, p53 (and Brca2) proteins.

**Table 2.** Fold changes in endogenous proteins after 15 min exposure to MG132

| Cell line | Rad51 | Brca2 | p53 |
| --- | --- | --- | --- |
| igm482 | $0.9 \pm 0.06$ | $1.5 \pm 0.07$ | $1.3 \pm 0.03$ |
| K133R-11 | $1.3 \pm 0.12^{\dagger}$ | $1.8 \pm 0.20^{\text{ns}}$ | $1.8 \pm 0.11^{\dagger}$ |
| K133A-6 | $2.0 \pm 0.09^{\dagger\dagger}$ | $1.7 \pm 0.07^{\text{ns}}$ | $1.9 \pm 0.15^{\dagger}$ |

(t-test, $^{\dagger\dagger}p \leq 0.01$ , $^{\dagger}p \leq 0.05$ , ns; not significant).

### Rad51 catalytic mutants feature elevated Rad51 polyubiquitination

To gain further information on the mechanism of proteasome-mediated degradation, we analyzed the extent of Rad51 polyubiquitination before and after MG132 treatment in the control igm482 hybridoma, as well in cell lines expressing the Rad51 catalytic mutants. As shown in Fig. 3a (lanes 2-7), various higher molecular weight Rad51 species are visible following immunoprecipitation with anti-Rad51 (H92) antibody and Western blotting with anti-Rad51 (H92) antibody. Further, Western blot analysis of these immunoprecipitates with anti-ubiquitin antibody detects two prominent ubiquitinated Rad51 species of ~50 kDa and ~85 kDa (lanes 11-16). Interestingly, compared to the untreated igm482 cells (lane 11), higher levels of the ubiquitinated Rad51 species are present in the untreated K133A-6 (lane 13) and in untreated K133R-11 immunoprecipitates (lane 15). Further, treating cells with MG132 increased the levels of these ubiquitinated-Rad51 species suggesting that Rad51 ubiquitination is transient (compare lanes 12, 14 and 16).

**Fig. 3.**
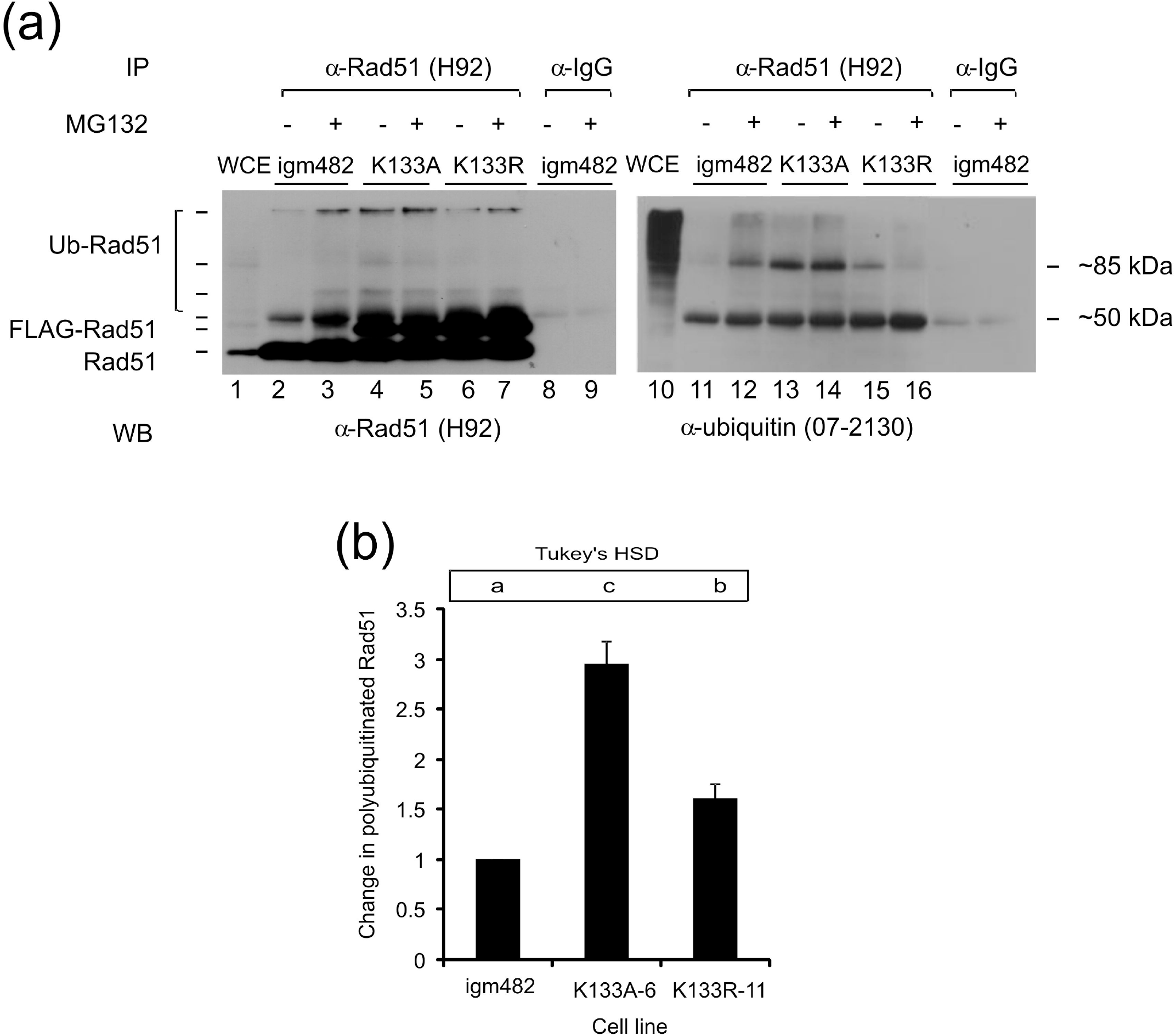
Rad51 polyubiquitination. (a) Analysis of Rad51 polyubiquitination. Whole cell extracts (WCE) from cells that were treated (+) with the proteasome inhibitor MG132 (10 μM) or untreated (-) were immunoprecipitated (IP) with anti-Rad51 (H92), followed by Western blot (WB) analysis with either the same anti-Rad51 (H92) antibody or with anti-ubiquitin antibody as indicated. Immunoprecipitations with non-specific rabbit IgG validate the specificity of the interaction. WCE from igm482 cells are included as a positive control for the immunoblot. (b) Fold change in Rad51 polyubiquitination compared to control igm482 cells ± standard error of the mean as determined by densitometric analysis of Western immunoblots from at least 3 independent experiments. One-way ANOVA and Tukey’s-HSD analysis were used to assess statistically significant differences between cell lines. Means indicated by the same lower-case letter are not statistically different (p≤0.05).

Next, we used densitometric analysis to quantify the extent of Rad51 poly-ubiquitination in the various cell lines. The amount of ubiquitinated Rad51 was first determined as a ratio of the amount of immunoprecipitated Rad51 using a serial dilution of the immunoprecipitations shown in Fig. 3A. We then normalized this data to the control igm482 hybridoma. These analyses were repeated for at least 3 independent experiments. As shown in Fig. 3b, the amount of polyubiquitinated Rad51 is ~3-fold higher in K133A-6 and ~1.5-fold higher in K133R-11 compared to the control igm482 cells supporting ubiquitin-mediated proteasome degradation.

### FLAG-tagged Rad51 proteins interact with endogenous Rad51, Brca2 and p53

Next, we sought to establish whether wild type FLAG-Rad51 and the Rad51-K133A and Rad51-K133R catalytic mutants were able to interact with endogenous Rad51, Brca2 and p53. To assess interactions with Rad51, extracted cellular protein was immunoprecipitated with anti-FLAG antibody (clone M2) and analyzed by Western blot with anti-Rad51 antibody (14B4). As shown in Fig. 4a, anti-FLAG antibody immunoprecipitates wild-type FLAG-Rad51 and endogenous wild-type Rad51 from hybridoma cell lines WT5 (lane 4) verifying wild-type FLAG-Rad51 association with endogenous Rad51. A similar association with endogenous Rad51 is observed for Rad51-K133A (lane 6) and Rad51-K133R (lane 8). Immunoprecipitations from control igm482 cells (lane 2) and with control IgG antiserum (lanes 9 and 10) validate the specificity of the assay.

**Fig. 4.**
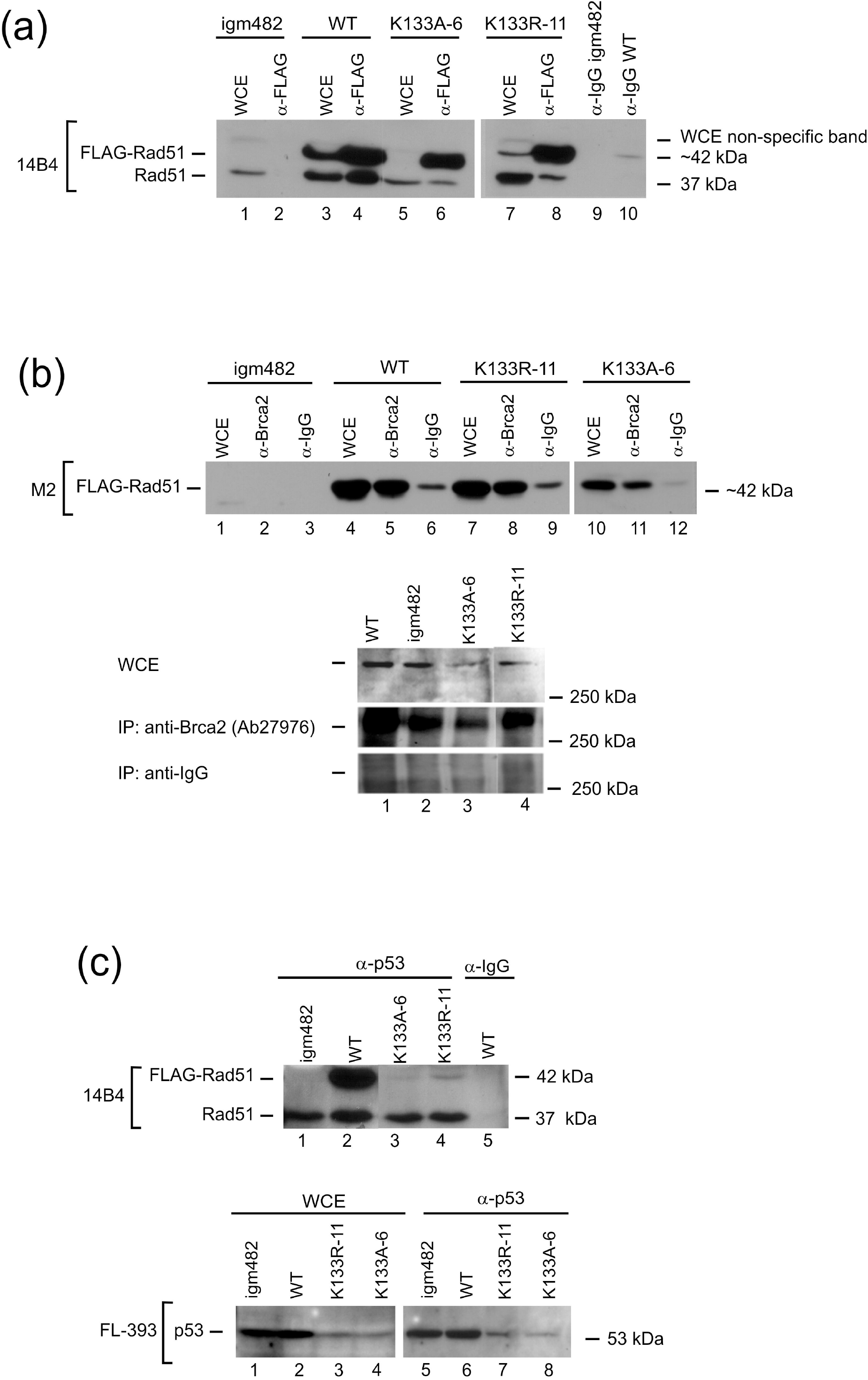
Interaction of catalytic Rad51 mutants with endogenous Rad51, Brca2 and p53. (a) Interaction between FLAG-tagged and endogenous Rad51. Protein complexes were immunoprecipitated with anti-FLAG antibody (M2) and analyzed by Western blot with anti-Rad51 antibody (14B4). Whole cell extracts (WCE) are included for comparison. Immunoprecipitations with non-specific rabbit IgG from control igm482 and cells expressing wild-type FLAG-tagged Rad51 were included to validate the specificity of the interaction. (b) Interaction between FLAG-tagged Rad51 and Brca2. Protein complexes were immunoprecipitated with either anti-Brca2 antibody (Ab27976) or non-specific rabbit IgG followed by Western blot analysis with anti-FLAG antibody (M2) or with anti-Brca2 (Ab27976) antibody. WCE are included for comparison. (c) Interaction between FLAG-Rad51, endogenous Rad51, and p53. Protein complexes were immunoprecipitated with anti-p53 (FL-393) and analyzed by Western blot analysis with anti-Rad51 antibody (14B4), or they were immunoprecipitated with anti-p53 (FL-393) antibody covalently bound to agarose beads followed by Western blot analysis with anti-p53 (FL-393) antibody. To validate the specificity of the assay, immunoprecipitations were also performed with non-specific anti-IgG.

Immunoprecipitations performed with anti-Brca2 antibody (Ab27976) followed by Western blot analysis with anti-FLAG antibody (clone M2)(Fig. 4b; panel 1) revealed that wild-type FLAG-Rad51 (lane 5), Rad51-K133R (lane 8) and Rad51-K133A (lane 11) interact with mouse Brca2. Immunoprecipitations performed with control igm482 cell extracts (lane 2) and the low level of non-specific FLAG-Rad51 binding by anti-IgG serum (lanes 3, 6, 9 and 12) support the specificity of the assay. To further validate the assay, we immunoprecipitated protein complexes with anti-Brca2 (Ab27976) and performed Western immunoblot analysis with the same antibody. As shown in Fig. 4b (panel 3; lanes 1-4), the antibody indeed immunoprecipitates Brca2. We also tested the anti-Brca2 (Ab27976) pulldowns for immunoprecipitation of endogenous Rad51 using anti-Rad51 antibody (14B4), but did not observe any reactive bands (data not shown). This was not surprising because our previous estimates suggest that the concentration of endogenous Brca2 is at least 4-fold lower than Rad51 requiring immunoprecipitations using at least twice the amount of whole cell extract (2 mg) and four-fold the amount of anti-Brca2 antibody (40 μg) to detect only a faint Rad51 signal equivalent to about 3% of the total endogenous Rad51.^26^

Immunoprecipitations performed with anti-p53 (FL-393) and analyzed by Western blot analysis with anti-Rad51 antibody (14B4) (Fig. 4c, panel 1) revealed that wild-type FLAG-Rad51 (lane 2) as well as Rad51-K133A (lane 3) and Rad51-K133R (lane 4) interact with mouse p53. Immunoprecipitations with anti-p53 in control igm482 cells (lane 1) and assessment of non-specific binding with anti-IgG antibody (lane 5) validate the specificity of the assay. To further validate the assay, we immunoprecipitated protein complexes with anti-p53 (FL-393)-conjugated agarose beads followed by Western blot analysis for p53 (anti-p53; FL-393). As shown in panel 2, the complexes precipitated by the antibody do contain p53 and the amounts immunoprecipitated from whole cell extracts of the catalytic mutants are reflective of the low level of p53 present in these cells.

Based on the above information, we conclude that FLAG-tagged wild-type Rad51 and the K133A and K133R catalytic Rad51 mutants can associate with endogenous Rad51, Brca2 and p53 *in vivo*.

### FLAG-tagged Rad51 proteins localize to the nucleus in response to DNA damage

To determine whether the Rad51 catalytic mutants retain the ability to enter the nucleus, we compared the cellular distribution of the proteins before and after cells were treated for 18 h with 600 nM DNA damaging agent mitomycin C (MMC) (Fig. 5). We observed nuclear entry of endogenous Rad51 in control igm482 cells (Fig. 5a; panel 1, compare lanes 6 and 9) and of wild-type FLAG-Rad51 and endogenous Rad51 in WT5 cells expressing wild type FLAG-Rad51 (Fig. 5b; panels 1 and 2, compare lanes 6 and 8). We also observed that the catalytic mutants Rad51-K133A (Fig. 5c; panel 1, compare lanes 6 and 8) and Rad51-K133R (Fig. 5d; panel 1, compare lanes 6 and 9) gain nuclear entry after treatment with MMC. The purity of the fractions was verified using anti-histone H1 antibody.

**Fig. 5.**
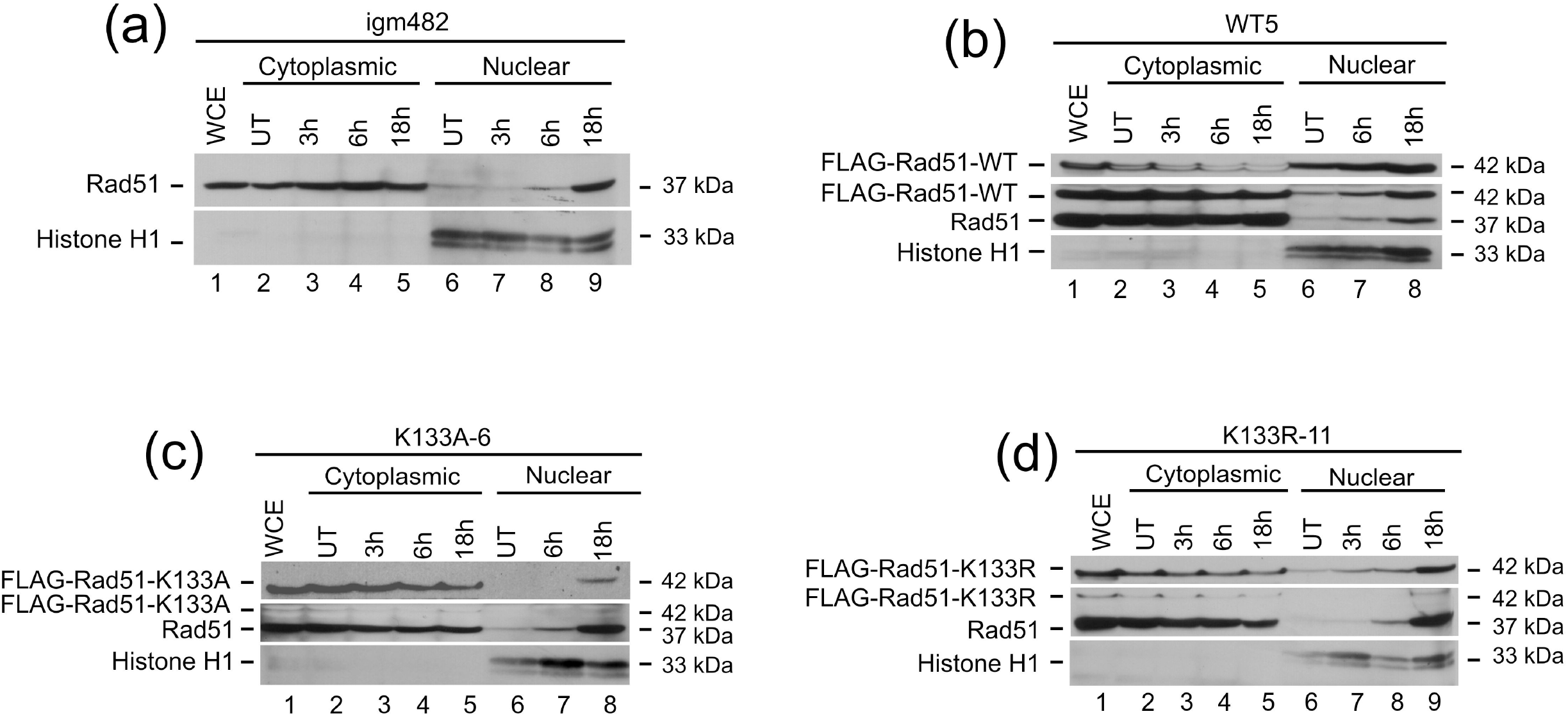
Nuclear translocation of Rad51. Western blot analysis was used to determine the cellular redistribution of Rad51 following MMC treatment of (a) igm482, (b) WT5, (c) K133A-6 and (d) K133R-11 cell lines. Cells were treated for 18 h with 600 nM MMC. The blots were probed with anti-Rad51 antibody (14B4) and/or anti-FLAG (clone M2) antibodies where appropriate. The purity of the fractions was verified using anti-histone H1 antibody. Abbreviations: WCE, whole cell extracts; UT, untreated; MMC, mitomycin C.

### Rad51 catalytic mutants inhibit the early steps of homologous recombination

We subsequently investigated whether the Rad51 catalytic mutants affected the new DNA synthesis that accompanies the early homology search and strand invasion steps of HR. As detailed in Si *et al*., recipient igm482 hybridoma cells are electroporated with a gene targeting vector (pTΔCμ_858/2290_) in which otherwise continuous homology (4.3 kb) to the single copy chromosomal immunoglobulin μ gene is interrupted by a double-strand gap (DSG) generating 858 bp and 2290 bp arms to the left and right of the DSG, respectively (Fig. 6a). Specific pairing between the homologous arms of the transferred vector and the cognate μ locus primes new DNA synthesis into the gapped region (3’ polymerization), which is subsequently detected by a sensitive PCR assay.

**Fig. 6.**
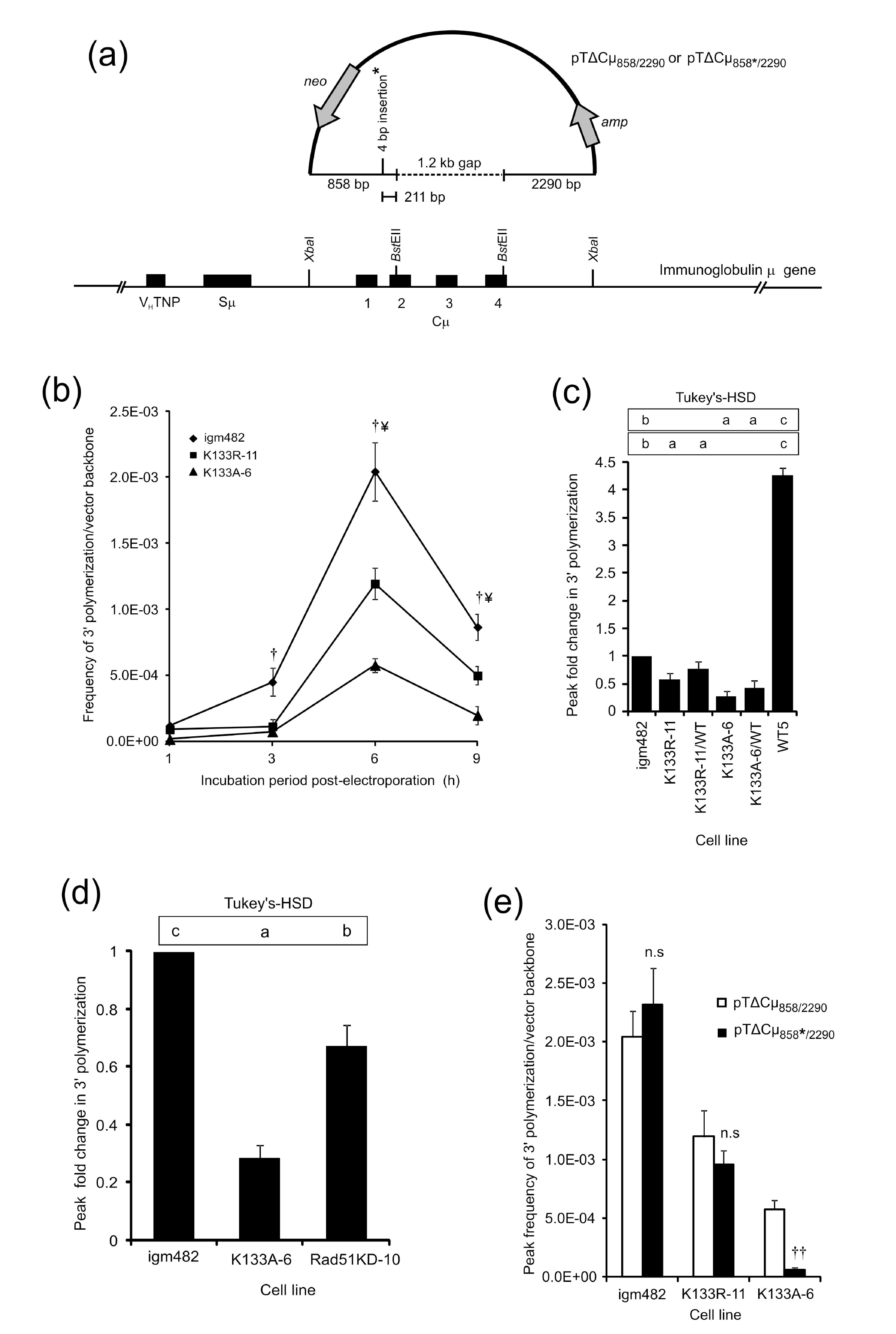
Effects of Rad51 catalytic mutants on early steps of HR (3’ polymerization). (a) Schematic diagram of the vectors pTΔCμ_858/2290_ and pTΔCμ_858*/2290_ used in the 3’ polymerization assay. In pTΔCμ_858*/2290_, otherwise continuous homology in the left vector arm is interrupted by a 4 bp insertion (positioned 211 bp from the vector 3’ end). The single-copy hybridoma chromosomal immunoglobulin μ gene serves as template for repair of the 1.2 kb vector-borne gap. (b) Kinetics of 3’ polymerization. The mean frequency of 3’ polymerization/vector backbone ± standard error of the mean is shown for each cell line and time point. The symbol (¥) indicates statistically significant differences between all three cell lines and daggers (f) indicate statistically significant differences from the control igm482 (*t*-tests, *p* ≤ 0.05). (c) Effect of wild-type Rad51 over-expression on 3’ polymerization in cell lines expressing Rad51 catalytic mutants. The mean peak fold change in 3’ polymerization (at 6 h) ± standard error of the mean relative to control igm482 cells is presented. The mean fold change in 3’ polymerization for the cell line WT5 expressing wild-type FLAG-Rad51 compared to igm482 was previously reported by Mundia *et al*.^23^ and is included here for comparison. Differences in 3’ polymerization between cell lines were determined by one-way ANOVA and Tukey’s-HSD analysis. Means indicated by the same lower-case letter are not statistically different (p≤0.05). (d) Comparison of 3’ polymerization in cell line K133A-6 and an igm482 derivative (Rad51KD-10) that is ~60% shRNA-reduced for Rad51^26^. The mean peak fold change in 3’ polymerization (at 6 h) ± standard error of the mean relative to control igm482 cells is presented. Differences in 3’ polymerization were determined by one-way ANOVA and Tukey’s-HSD analysis. Means indicated by the same lower-case letter are not statistically different (p≤0.05). (e) Effect of heterology on 3’ polymerization in Rad51 catalytic mutants. The peak frequencies of 3’ polymerization/vector backbone for control igm482, K133R-11 and K133A-6 cell lines ± standard error of the mean, following transfection with homologous (pTΔCμ_858/2290_) and heterologous (pTΔCμ_858*/2290_) vectors. Differences in 3’ polymerization between homologous (pTΔCμ_858/2290_) and heterologous (pTΔCμ_858*/2290_) vectors were determined by unpaired *t*-tests. †† p=0.0012. n.s; not significant.

We measured 3’ polymerization in control igm482 hybridoma cells and in cell lines K133R-11 and K133A-6. While the kinetics of 3’ polymerization are not altered, the Rad51 catalytic mutants cause a significant decrease in the efficiency of 3’ polymerization relative to control igm482 cells at 3, 6, and 9 h (Fig. 6b). Compared to control igm482 cells, the reduction in the peak frequencies of 3’ polymerization (at 6 h) is ~70% for cell line K133A-6 and ~40% for cell line K133R-11.

It was of interest to determine whether the significant deficiency in 3’ polymerization caused by expression of the Rad51 catalytic mutants, K133A and K133R could be reversed by provision of extra wild-type Rad51. Consequently, we measured 3’ polymerization in K133R-11 and K133A-6, along with derivatives (K133R-11/WT and K133A-6/WT) expressing an excess of wild-type FLAG-Rad51 relative to β-actin (Fig. 1a top panel, compare lanes 1, 5 and 9). Notably, as shown in Fig. 6c, provision of excess wild-type FLAG-Rad51 slightly improves 3’ polymerization efficiency but does not completely rescue the 3’ polymerization defect caused by the mutant Rad51-K133A and Rad51-K133R proteins, and does not achieve the level of 3’ polymerization stimulation observed in WT5 (~4-fold^23^). Therefore, although wild-type FLAG-Rad51 stimulates 3’ polymerization in the background of endogenous Rad51, it cannot do so when co-expressed with the Rad51-K133A or Rad51-K133R mutant proteins.

We subsequently wondered whether the reduction in 3’ polymerization seen in cells expressing the catalytic mutants (Fig. 6b,c) was simply due to the reduction of Rad51 (and Brca2) observed in Fig. 1a rather than a mechanistic effect of the mutants on HR. To test this, we compared the 3’ polymerization in K133A-6 with a Rad51-knockdown cell line (Rad51KD-10). This cell line is ~60% reduced for Rad51 via shRNA-mediated depletion and shows a concomitant reduction in Brca2 and p53 via a cellular rebalancing mechanism.^26^ Since K133A-6 has approximately the same level of endogenous Rad51 as Rad51KD-10, we hypothesized that the efficiency of 3’ polymerization would be similar in K133A-6 and Rad51KD-10 if the effect were solely due to the reduction in protein levels. As shown in Fig. 6d, K133A-6 is significantly more defective in 3’ polymerization than Rad51KD-10, suggesting that an additional mechanism of HR inhibition is at play in cells expressing Rad51 catalytic mutants.

We hypothesized that the Rad51-K133A and Rad51-K133R catalytic mutants inhibit 3’ polymerization (as well as gene targeting^14^) by interfering with Rad51 nucleoprotein filament function. To investigate this, we examined whether the ATP-binding deficient K133A protein (in cell line K133A-6) and the ATP-hydrolysis deficient K133R protein (in cell line K133R-11) were altered in their capacity to form heteroduplex DNA (hDNA) during 3’ polymerization. We exploited the vector pTΔCμ_858*/2290_, a pTΔCμ_858/2290_ derivative in which the otherwise continuous 858 bp of μ gene homology in the left vector arm is interrupted by a 4 bp insertion (Fig. 6a). During gene targeting, hDNA can form over distances >1 kb from invading 3’ vector ends ^24,34^ suggesting the likelihood of it spanning the 4 bp insertion positioned only 211 bp from the 3’ end of the transferred pTΔCμ_858*/2290_ vector.

At peak frequency in control igm482 cells, 3’ polymerization measured using the pTΔCμ_858*/2290_ vector is not significantly different from that observed with vector pTΔCμ_858/2290_ bearing the wild-type μ sequence (Fig. 6e) suggesting that in the presence of endogenous wild-type Rad51, the 4 bp mismatch does not affect the efficiency of hDNA formation. In cell line K133R-11, there are also no significant differences between the vectors; while the overall efficiency of 3’ polymerization is reduced relative to igm482, the capacity to tolerate hDNA was not negatively impacted. In marked contrast, in cell line K133A-6, the 4 bp mismatch in pTΔCμ_858*/2290_ significantly impairs 3’ polymerization relative to the wild-type μ sequence in pTΔCμ_858/2290_ supporting a strong requirement for Rad51 ATP binding in hDNA formation. Therefore, our results suggest that the Rad51-K133R and Rad51-K133A proteins both display dominant-negative effects with respect to Rad51 nucleoprotein filament function during the early steps of homologous recombination. Further, the dominant-negative effects can be separated, with Rad51-K133A displaying a strong tendency to disrupt 3’ polymerization in the presence of DNA sequence heterology.

## DISCUSSION

The focus of this study was the evaluation of Rad51 catalytic mutant alleles on the Rad51 nucleoprotein filament *in vivo*. Accordingly, we utilized mouse hybridoma cell lines stably expressing ectopic N-terminal FLAG-tagged wild-type Rad51, or the FLAG-tagged Rad51-K133A and Rad51-K133R catalytic mutants deficient in ATP binding and ATP hydrolysis, respectively.^8–10,12^’ Intrinsic to proper Rad51 nucleoprotein filament formation/function is the maintenance of normal endogenous levels of homologous recombination proteins. Previously, we showed that levels of endogenous Rad51 and Brca2 are dependent on each other and when reduced, are associated with a lower level of endogenous p53, presumably to permit continued cell growth.^26^ Therefore, we questioned whether ectopic expression of the Rad51 catalytic mutants affected the endogenous levels of these critical homologous recombination proteins. Quite unexpectedly, endogenous levels of Rad51, Brca2 and p53 proteins were significantly lower in cell lines expressing Rad51-K133A and Rad51-K133R. Thus, this study further supports a cellular mechanism that permits re-adjustment of the levels of endogenous Rad51, Brca2 and p53 likely to facilitate HR and cell survival.

As a first step in understanding the mechanism by which these critical homologous recombination protein levels are altered, RT-PCR analysis of *Rad51, Brca2* and *p53* transcript levels was performed. This analysis revealed that the alteration in the levels of these proteins was not accounted for by changes in mRNA levels. Comparison of protein levels in control igm482 cells and cells expressing the catalytic mutant Rad51 treated with an inhibitor of protein biosynthesis, cycloheximide^28^ or the proteasome-inhibitor, MG132^21^, suggests expression of the catalytic mutants enhances the susceptibility of the endogenous Rad51, Brca2, and p53 proteins to proteasome-mediated degradation. Indeed, we obtained evidence for increased Rad51 polyubiquitination in cells expressing the Rad51 catalytic mutants.

The control of HR represents a highly regulated balance between events that activate proteins to promote recombination and ensure genome integrity, and those that appropriately down-regulate HR. Indeed, a link between the proteasome and the control of HR has been identified. The DSS1 protein (equivalent to Sem1 in *S. cerevisiae*^35^ interacts with BRCA2 and is important for its stability.^36,37^ Moreover, both DSS1 and BRCA2 interact with two important proteasome components, RPN3 and RPN7, and it is possible that BRCA2 might localize the proteasome to sites of DSB repair.^38^ Additionally, yeast Sem1 is recruited along with 19S and 20S proteasome subcomplexes to HO-induced double-strand breaks (DSB) *in vivo*.^39^ Further, Rad51 regulation is by ubiquitin-mediated proteasome degradation; Rad51 is a substrate of numerous ubiquitin ligases including UBC9 protein, which is homologous to the ubiquitin-conjugating enzymes Hus5 of fission yeast and Ubc9 of budding yeast, and this interaction targets Rad51 for cytosolic degradation.^40^ Rad51 is also a target of F-box DNA helicase 1 (FBH1) which acts as part of the SCF (SKP1, CUL1 and F-box) complex.^41,42^ *In vitro,* the SCF^FBH1^ complex has ubiquitin-ligase (E3) activity and monoubiquitylates multiple lysine residues of Rad51.^42^ In yeast, fbh1-mediated ubiquitination regulates the levels of Rad51 protein.^41^ Thus, altogether, the above results provide additional support for the importance of the ubiquitin-proteasome pathway in regulating the levels of endogenous Rad51, Brca2 and p53 in response to ectopic expression of the Rad51-K133A and Rad51-K133R catalytic mutants.

In mammalian cells, the recombination mediator BRCA2 regulates formation of the RAD51 nucleoprotein filament through eight BRC repeats that bind RAD51 monomers selectively loading them onto 3’ ending, single-stranded DNA overhangs.^4–6^ Thus, in considering Rad51 nucleoprotein filament formation, an important question was whether the Rad51 catalytic mutants were able to interact with endogenous Brca2 and Rad51. Our specific immunoprecipitation studies reveal that FLAG-tagged wild-type Rad51, as well as the FLAG-tagged Rad51-K133A and Rad51-K133R catalytic mutants interact with endogenous Rad51 and Brca2 (as well as p53) proteins *in vivo*. These findings extend previous studies reporting specific complex formation between the wild-type versions of these proteins^26,27,43,44^ and interactions between GFP-tagged human RAD51-K133R and -K133A proteins, and endogenous human RAD51, BRCA2 BRC3 repeat, ssDNA and ATP^10^. Rad51-K133A association with Brca2 implies that interactions with Brca2 do not require ATP-bound Rad51, a notion supported by previous structure/function studies of BRCA2 BRC repeat: RAD51 interactions^45^ and studies with full-length purified BRCA2^5^. Therefore, the observed interaction of endogenous Rad51 and Brca2 with FLAG-tagged wild-type Rad51, and the Rad51-K133A and Rad51-K133R catalytic mutants provides the prerequisite interactions necessary for homotypic Rad51 nucleoprotein filament formation [similar to that observed for *S. cerevisiae* between wild-type Rad51 and Rad51-K191A (the yeast equivalent to mammalian Rad51-K133A)]^11^.

From the findings presented above, we were prompted to investigate nucleoprotein filament function by studying the new DNA synthesis (3’ polymerization) that accompanies the early homology search and strand invasion steps of HR *in vivo*.^22^ Compared to the basal level of 3’ polymerization in control igm482 cells and the elevation in 3’ polymerization observed in WT5 expressing wild-type FLAG-Rad51^14,23^, cell lines expressing the Rad51-K133A and Rad51-K133R catalytic mutants are significantly defective in 3’ polymerization. Interestingly, the inhibitory effects of the Rad51 catalytic mutants are not equivalent; Rad51-K133A is considerably more toxic. Importantly, 3’ polymerization in cell lines expressing Rad51-K133A or Rad51-K133R was not restored following provision of an additional ~ 2-fold excess of ectopic wild-type FLAG-Rad51 (a level similar to that previously found to elevate 3’ polymerization in the background of endogenous wild-type Rad51)^23^. Therefore, we are the first to show that the toxic effects of the Rad51 catalytic mutants on HR arise via dominant-negative inhibition of wild-type Rad51 function and occur at the fundamental level of the Rad51 nucleoprotein filament *in vivo*.

A basic question relates to the mechanism by which the Rad51-K133R and Rad51-K133A catalytic mutants exert their dominant-negative effects on the Rad51 nucleoprotein filament, that is, whether the reduction in 3’ polymerization is the result of reduced formation of the Rad51 nucleoprotein filament or conversely, a decrease in its activity. We were guided in our approach to this question by several previous observations in microbial systems, where wild-type RecA, but not RecA-K72R (similar to Rad51-K133R) or non-hydrolysable ATP analogs (ATP-γ-S) promoted efficient strand exchange between heterologous DNA substrates.^46–54^ To investigate whether 3’ polymerization was affected by DNA sequence heterology and if the Rad51 catalytic mutants differed in their response, we modified our usual 3’ polymerization assay to include a derivative in which homology is interrupted by a 4 bp insertion. Our results show that in control igm482 cells, the frequency and kinetics of 3’ polymerization were similar for homologous and heterologous DNA substrates. Thus, nucleoprotein filaments active in 3’ polymerization and formed with endogenous wild-type Rad51 are of similar effectiveness regardless of DNA heterology. In contrast, the efficiency of 3’ polymerization is significantly reduced and by a similar extent, for homologous and heterologous plasmid substrates in cells expressing Rad51-K133R. The deficiency in 3’ polymerization for both plasmid substrates might be accounted for by Rad51-K133R interfering (or competing) with endogenous Rad51 essentially crippling nucleoprotein filament formation. However, given our observed interactions between wild-type Rad51 and Rad51-K133R (along with similar interactions between endogenous Rad51 and Rad51-K191A in *S. cerevisiae*.^11^), the possibility of homotypic Rad51 associations appears likely. Accordingly, our data suggest that compared to wild-type Rad51, a mixed Rad51:Rad51-K133R nucleoprotein filament has similar, but significantly reduced activity in both strand exchange and the capacity to tolerate DNA heterology. Like Rad51-K133R, deficiency in 3’ polymerization in cells expressing Rad51-K133A might result from reduced Rad51 nucleoprotein filament formation (either through interference or competition with wild-type Rad51). However, support for a homotypic Rad51:Rad51-K133A nucleoprotein filament is provided by our findings of interactions between endogenous Rad51 and Rad51-K133A, as well as reported interactions between Rad51-K133A, and endogenous Brca2 and Xrcc3.^10^ Further, even though Rad51-K133A has an approximately 100-fold reduced affinity for ATP binding, Rad51-depleted cell lines expressing Rad51-K133A are still able to function in the repair of DNA DSB.^10^ Perhaps the most compelling evidence for a mixed Rad51:Rad51-K133A nucleoprotein filament is the different 3’ polymerization frequencies for homologous and heterologous plasmid substrates: compared to control cells and those expressing Rad51-K133R, expression of Rad51-K133A results in a significant crippling of 3’ polymerization when homologous plasmid DNA is used and an even more significant impairment under conditions where strand exchange requires tolerance of sequence heterology.

The above information suggests an interesting mechanistic dichotomy in the way Rad51-K133R and Rad51-K133A proteins exert dominant-negative effects on 3’ polymerization. As indicated above, Rad51-K133R binds ATP, but is deficient in ATP-hydrolysis.^8–10, 12^ ATP-binding induces a conformational change in Rad51 that favors strong ssDNA-RAD51 interactions and therefore, the ATP-bound Rad51-nucleoprotein filament remains stable and active ^9^; *in vitro,* free ATP in solution stabilizes yeast Rad51 nucleoprotein filaments five-fold compared to ATP-free buffer.^55^ However, ATP hydrolysis is required for the sinuous removal of Rad51-ADP and consequently, filaments formed with Rad51-K133R are defective for Rad51 turnover, their stability is altered ^56,57^ and homologous recombination is affected.^9,12^ Similarly, in *S. cerevisiae,* Rad51-K191R (similar to mouse Rad51-K133R) forms hyperstable Rad51 nucleoprotein filaments that are deficient in HR.^58,59^ Given the ATP hydrolysis deficiency, defective Rad51 turnover would likely reduce the amount of free Rad51 available and hinder timely access by DNA polymerase, thus impairing 3’ polymerization. Interestingly, our studies show that the inhibitory effects of Rad51-K133R on 3’ polymerization are not alleviated by expression of excess ectopic wild-type Rad51 suggesting that Rad51-K133R:wild-type Rad51 associations are inherently non-productive and drive the dominant-negative HR phenotype.

In the case of the Rad51-binding deficient Rad51-K133A mutant^8–10,12^, the stark difference in 3’ polymerization between homologous and heterologous plasmid substrates supports formation of a homotypic Rad51:Rad51-K133A nucleoprotein filament, but one which is severely defective in strand invasion/exchange and heterology by-pass. The significantly reduced ATP-binding ability of Rad51-K133A likely impacts Rad51 nucleoprotein filament activity, especially its capacity to drive strand exchange through regions of heterology. This could occur through formation of short, less active Rad51:Rad51-K133A multimers, or perhaps, a more extended Rad51 nucleoprotein filament that displays reduced activity for strand exchange and/or heterology by-pass.

Given the results of the present study which show that Rad51-K133A and Rad51-K133R associate with endogenous Rad51 and Brca2 (and p53) and mediate negative effects with respect to Rad51 nucleoprotein filament formation and/or function in strand exchange and heterology by-pass, it seems highly likely that toxic Rad51-nucleoprotein filaments and associated HR deficits could in turn trigger the observed proteasome-mediated destruction of key players, such as, Rad51 and Brca2 which further exacerbates the HR defect. If defective nucleoprotein filaments are targeted for degradation, endogenous Rad51 is depleted, which triggers Brca2 depletion to maintain the balance of the two proteins in the cell. The ensuing reduction of HR function may explain our observation of lowered p53 protein as a means of promoting cell survival, similar to that observed in knockouts of Rad51^60^, Brca1, Brca2^61,62^, knockdowns of Rad51 and Brca2^26^, and in human BRCA1-and BRCA2-associated breast tumors^63,64^. Since Rad51 is also important in restarting stalled replication forks by HR, and both Rad51-K133A and Rad51-K133R are inhibitory to this process^18^, it is expected that negative effects would be manifested genome-wide providing an explanation for the observed toxic phenotypic effects of the Rad51 catalytic mutants such as increased sensitivity to DNA-damaging agents and impaired growth.^14,15,18^

In conclusion, we utilized cell lines stably expressing wild-type FLAG-Rad51, Rad51-K133A and Rad51-K133R in order to study the role of ATP binding and hydrolysis during the early homology search and strand invasion events of HR *in vivo*. The results of this study show that even low levels of the Rad51-K133A and Rad51-K133R catalytic mutants affect the formation and/or function of the Rad51 nucleoprotein filament, and reveal that they do so in a dichotomous way: Rad51-K133R in concert with endogenous Rad51 presumably forms Rad51-nucleoprotein filaments, but the ATP hydrolysis deficiency of K133R likely renders such filaments defective for Rad51 turnover explaining the deficit in 3’ polymerization. In contrast, the ATP binding defective Rad51-K133A protein likely compromises Rad51 nucleoprotein filament formation and/or its activity as evidenced by its large 3’ polymerization defect and significantly reduced capacity to drive strand exchange through regions of heterology. Expression of the catalytic mutant Rad51-K133R and Rad51-K133A (but not wild-type Rad51) proteins is associated with accelerated proteasome-mediated depletion of endogenous Rad51 and Brca2, and the ensuing deficit in HR presumably selects for low p53 as an aid in promoting cell survival. Importantly, we show that the toxic effects of Rad51-K133R and Rad51-K133A are manifested in the background of endogenous wild-type Rad51 and significantly, are not reversed by provision of excess ectopic wild-type Rad51. The perturbation of wild-type Rad51 function by expression of even low levels of the Rad51-K133R and Rad51-K133A catalytic mutants highlights their dominant-negative effects with respect to Rad51 nucleoprotein filament function during HR.

## ACKNOWLEDGEMENTS

We acknowledge Vatsal Desai for the construction of the pTΔCμ_858*/2290_ vector. This work was supported by an Operating Grant from the Canadian Institutes of Health Research (CIHR) to M.D.B and University of Guelph international graduate scholarships to M.M.M.

---

**Fig. S1. Elevated endogenous Rad51 levels in cell lines overexpressing wild-type Rad51 are due to multiple opportunities for internal translation initiation.** (a) Nucleotide sequence alignment of an N-terminal segment of FLAG-tagged wild-type mouse Rad51 (GenBank accession number NM_011234) and FLAG-Rad51 in which the three potential internal ATG (AUG) translation initiation codons have been replaced with AGG. (b) Western blot analysis of endogenous and FLAG-Rad51 expression in WT5 and three representative N-terminal mutant cell lines. β-actin is presented as a control for protein loading. The three amino acid substitutions are predicted to slightly increase the molecular weight of the FLAG-tagged Rad51 and therefore, the faster migration of the N-terminal FLAG-tagged Rad51 in the mutants is an anomaly. We attributed this difference to “gel-shifting”: a phenomenon that is widely reported where amino acid changes unpredictably alter the electrophoretic mobility of a given protein during sodium dodecyl sulphate polyacrylamide gel electrophoresis.^65–68^

